# rPanglaoDB: an R package to download and merge labeled single-cell RNA-seq data from the PanglaoDB database

**DOI:** 10.1101/2021.05.28.446161

**Authors:** Daniel Osorio, Marieke L. Kuijjer, James J. Cai

## Abstract

**Motivation:** Characterizing cells with rare molecular phenotypes is one of the promises of high throughput single-cell RNA sequencing (scRNA-seq) techniques. However, collecting enough cells with the desired molecular phenotype in a single experiment is challenging, requiring several samples preprocessing steps to filter and collect the desired cells experimentally before sequencing. Data integration of multiple public single-cell experiments stands as a solution for this problem, allowing the collection of enough cells exhibiting the desired molecular signatures. By increasing the sample size of the desired cell type, this approach enables a robust cell type transcriptome characterization.

**Results:** Here, we introduce rPanglaoDB, an R package to download and merge the uniformly processed and annotated scRNA-seq data provided by the PanglaoDB database. To show the potential of rPanglaoDB for collecting rare cell types by integrating multiple public datasets, we present a biological application collecting and characterizing a set of 157 fibrocytes. Fibrocytes are a rare monocyte-derived cell type, that exhibits both the inflammatory features of macrophages and the tissue remodeling properties of fibroblasts. This constitutes the first fibrocytes’ unbiased transcriptome profile report. We compared the transcriptomic profile of the fibrocytes against the fibroblasts collected from the same tissue samples and confirm their associated relationship with healing processes in tissue damage and infection through the activation of the prostaglandin biosynthesis and regulation pathway.

**Availability and Implementation:** rPanglaoDB is implemented as an R package available through the CRAN repositories https://CRAN.R-project.org/package=rPanglaoDB.

**Supplementary information:** Code to replicate the case example and figure 1 is available at https://github.com/dosorio/rPanglaoDB

## 1. Introduction

Merging and integrating the count matrices derived from multiple independent public single-cell RNA sequencing (scRNA-seq) experiments allows for better evaluation of the biological patterns in cell composition of tissues, as well as the identification of patterns of gene expression and gene regulation that are consistent across cells of the same cell type obtained from independent samples (Swamy, et al., 2021). A good quality integration of multiple datasets allowing comparison and contrasting of the data across different projects begins with a consistent preprocessing of the samples, using the same reference genome and the same quantification method to generate the count matrices (Lachmann, et al., 2018). These steps are then followed by the batch effect removal, normalization, cell type annotation, and characterization of samples and cell-types (Butler, et al., 2018; Hie, et al., 2019; Korsunsky, et al., 2019; Luecken, et al., 2020; Stuart, et al., 2019).

PanglaoDB is a secondary scRNA-seq database that reports the annotated count matrices for thousands of human and mice scRNA-seq experiments deposited in the sequence read archive (SRA) database of the National Center for Biotechnology Information (NCBI). Samples available in the PanglaoDB database are uniformly processed with the ‘alona’ package and made available in a web based unified framework at https://panglaodb.se/ (Franzen and Bjorkegren, 2020; Franzen, et al., 2019). However, the PanglaoDB database reports each sample on a single web page and does not offer options to automatically download or merge multiple available datasets based on molecular phenotypes or specific cell-type composition of the samples.

For that reason, here, we introduce rPanglaoDB, an R package to download and merge the uniformly processed and annotated scRNA-seq data provided by the PanglaoDB database. The package contains a comprehensive set of functions for filtering samples by organism and tissue from which the cells were collected, for cell type and signature marker genes expressed by the cells, as well as for the quality control and merging of the downloaded datasets. The final output of rPanglaoDB is a Seurat object, facilitating the downstream analysis and characterization of the data.

## 2. Material and Methods

rPanglaoDB includes four main functions, two for querying the list of samples available in the database, one to query the cell-type composition of the samples and one to download the samples’ count matrices and associated annotations.

### Querying samples in the database

Currently available samples can be accessed through two functions: ‘getSampleList’ which returns the list of all samples included in the database together with their associated annotations, such as the SRA database identifiers, the species and tissue from which the cells were collected, the protocol used, and the number of cells included in the sample, and ‘getMarkers’ which returns the list of clusters of cells in which a user-defined set of markers is expressed.

### Querying samples’ composition information from the database

Cell-type and number of cells by cell-type for the samples included in the database can be accessed through the ‘getSampleComposition’ function. It allows filtering of the results by the metadata associated to the sample returned by the ‘getSampleList’ function.

### Downloading samples’ count matrices and associated annotations

Once identified the samples, as well as the count matrices and the associated annotations can be downloaded using the ‘downloadSamples’ function. This function includes an option to return each sample as an independent Seurat object or to merge all samples into one Seurat object.

## 3. Results

Using the ‘getMarkers’ function, we identified a cluster of cells in the SRS3121028 sample derived from skin wound tissues (3 days after scab detachment) expressing *CD34, ACTA2, FN1*, Collagen V, *FAP, SIRPA* and the lack of expression of *CSF1R* (Fig. 1-A). Such combinatorial expression of genes differentiate fibrocytes from macrophages and fibroblasts (Lim, et al., 2018; Pilling, et al., 2009; Reilkoff, et al., 2011). Fibrocytes are associated with fibrosis, autoimmunity, cardiovascular disease and asthma, among other pathologies (Reilkoff, et al., 2011). Since, fibrocytes are marrow-derived cells that differentiate into fibroblasts-like phenotypes, they are usually wrongly labeled as fibroblasts. Thus, using the ‘getSamples’ function in rPanglaoDB, we downloaded all fibroblasts available from dermis samples in the database (SRA accessions: SRS3121028 and SRS3121030) (Lim, et al., 2018). We merged a total of 2,172 cells and processed the associated scRNA-Seq data using the Seurat package recommended pipeline (Stuart, et al., 2019). Datasets were integrated using Harmony and further corroboration of the marker genes defining their identity as fibrocytes (*CD34, ACTA2, COL5A1, COL5A2, COL5A3, FN1, FAP, SIRPA, PTPRC, MME*, and *SEMA7A*) was assessed using the Nebulosa package (Fig. 1-B) (Alquicira-Hernandez and Powell, 2021; Korsunsky, et al., 2019; Reilkoff, et al., 2011). Differential expression analysis of the cluster 8, which contained 157 fibrocytes (Fig. 1-A), against all the fibroblasts in the samples was performed using the MAST package (Fig. 1-C); returning 50 upregulated genes in fibrocytes associated with the *TGF-beta regulation of extracellular matrix, ECM-receptor interaction, Prostaglandin biosynthesis and regulation, Notch signaling pathway, Interleukin-5 regulation of apoptosis, Integrin beta-5 pathway, Oncostatin M, Hematopoietic cell lineage* and *Inflammatory response pathway* (FDR < 0.05using the hypergeometric-test through the enrichR package) (Finak, et al., 2015; Xie, et al., 2021). We cross validated the enrichment of the genes associated with the *Prostaglandin biosynthesis and regulation* pathway (*ANXA3, S100A10, ANXA5, ANXA2, PTGIS, ANXA1, S100A6, PTGS1*, and *HPGD*) using the Gene Set Enrichment Analysis (GSEA) approach included in the fgsea package (FDR = 0.01, Fig. 1-D) and the single-sample Gene Set Enrichment Analysis (ssGSEA) included in the GVSA package (FDR = 2.18 × 10^−18^, Fig. 1-E) (Hanzelmann, et al., 2013; Korotkevich, et al., 2021). All other associations did not pass the FDR threshold in all the other approaches applied (GSEA and ssGSEA). Our results show the potential of rPanglaoDB as a tool to collect cells with rare molecular phenotypes from the PanglaoDB database. We anticipate its use in the construction of atlases to characterize the different molecular phenotypes exhibited by different cell types in different tissues and organisms. We also provide the first unbiased, highly specific characterization of the fibrocyte transcriptome in skin wound tissues and confirm their association with healing processes in tissue damage and infection through the activation of the *Prostaglandin Biosynthesis and Regulation Pathway* (Grieb, et al., 2011; Zhang, et al., 2018).

**Figure 1.**
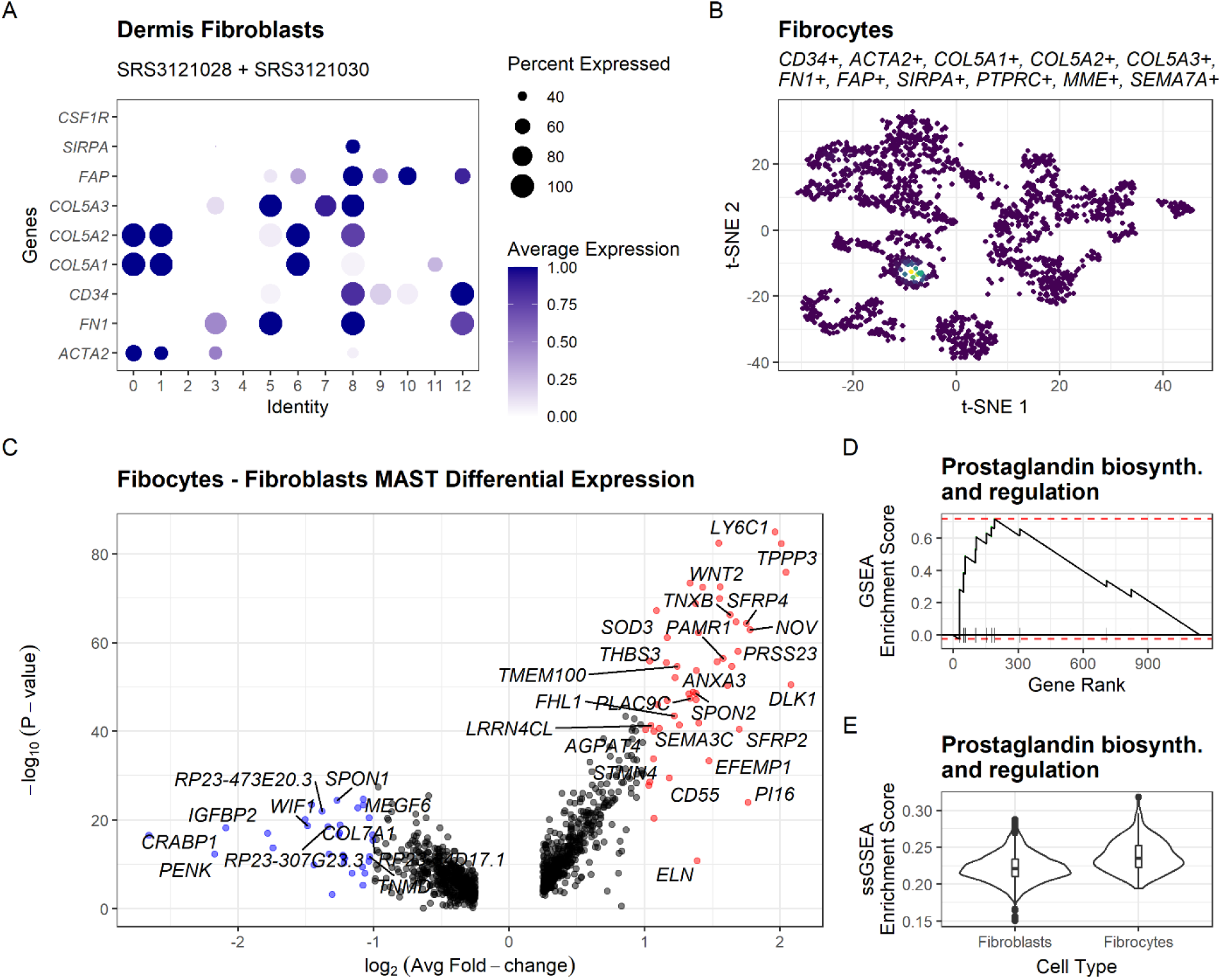
Characterization of the fibrocytes transcriptome. **(A)** Identification of cells expressing marker genes that differentiate fibrocytes’ identity from macrophages and fibroblasts (*CD34, ACTA2, FN1, Collagen V, FAP* and *SIRPA)*. **(B)** Cross validation of the identified cells expressing *CD34*^+^, *ACTA2*^+^, *COL5A1*^+^, *COL5A2*^+^, *COL5A3*^+^, *FN1*^+^, *FAP*^+^, *SIRPA*^+^, *PTPRC*^+^, *MME*^+^, and *SEMA7A*^+^ by kernel density estimation through the Nebulosa package. **(C)** Volcano plot displaying the differential expression between fibrocytes and the fibroblasts collected in the same merged samples. **(D)** Enrichment of the *prostaglandin biosynthesis and regulation pathway* using GSEA through the fgsea package. **(E)** Enrichment of the *prostaglandin biosynthesis and regulation pathway* using ssGSEA through the GSVA package.

## 4. Acknowledgments

D.O. was supported by the Marie Skł odowska-Curie Postdoctoral Scientia Fellows program from the Faculty of Medicine at the University of Oslo, co-funded by the EU Horizon 2020 MSCA COFUND. M.L.K was supported by the Norwegian Research Council, Helse Sør-Ø st, and University of Oslo through the Centre for Molecular Medicine Norway (NCMM).

